# White matter neuroanatomical predictors of aphasic verb retrieval

**DOI:** 10.1101/2020.09.30.316844

**Authors:** Haley C. Dresang, William D. Hula, Fang-Cheng Yeh, Tessa Warren, Michael Walsh Dickey

## Abstract

**Background:** Current neurocognitive models of language function have been primarily built from evidence regarding object naming, and their hypothesized white matter circuit mechanisms tend to be coarse-grained.

**Methods:** In this cross-sectional, observational study, we used novel correlational tractography to assess the white matter circuit mechanism behind *verb retrieval*, measured via action picture-naming performance in adults with chronic aphasia.

**Results:** The analysis identified tracts implicated in current neurocognitive dual-stream models of language function, including the left inferior fronto-occipital fasciculus, inferior longitudinal fasciculus, and arcuate fasciculus. However, the majority of tracts associated with verb retrieval were not ones included in dual-stream models of language function. Instead, they were projection pathways that connect frontal and parietal cortices to subcortical regions associated with motor functions, including the left corticothalamic pathway, frontopontine tract, parietopontine tract, corticostriatal pathway, and corticospinal tract.

**Conclusions:** These results highlight that cortico-subcortical projection pathways implicated in motor functions may be importantly related to language function. This finding is consistent with grounded accounts of cognition and may furthermore inform neurocognitive models.

**Impact Statement:** This study suggests that in addition to traditional dual-stream language fiber tracts, the integrity of projection pathways that connect frontal and parietal cortices to subcortical motor regions may be critically associated with verb-retrieval impairments in adults with aphasia. This finding challenges neurological models of language function.

## 1. Introduction

Prevailing neurolinguistic models are based on the characterization of cortical and subcortical substrates supporting object naming and noun retrieval. But there is more to language than nouns. Verbs engage distinct types of knowledge from nouns, serve different functions in language (Vigliocco *et al.*, 2011 for review), and can be differentially impaired in aphasia (Caramazza and Hillis, 1991; Miceli *et al.*, 1988). Verb processing is also important clinically. It is both a common locus of impairment in aphasia (Mätzig *et al.*, 2009) and a central focus of efficacious language treatments (Loverso *et al.*, 1987; Edmonds, 2016). A better understanding of the neural correlates of action naming and verb retrieval could broaden the scope of our neurolinguistic models as well as improve our understanding of which individuals with aphasia might be most likely to achieve positive outcomes from verb-based treatments. It could also be relevant to competing models of human cognition that disagree regarding the contribution of the motor system to language processing.

Although many functional neuroimaging and lesion studies have examined action words and concepts, we still know relatively little about the specific white matter tracts associated with verb retrieval. Many neuroimaging studies of verbs that report white matter correlates do not examine or identify specific fiber tracts. And although lesion-deficit approaches can characterize white matter damage correlated with behavioral impairments, studies using these approaches seldom describe *specific tracts* that are associated with impaired verb retrieval. The current study begins to remedy this by identifying specific white matter tracts associated with verb retrieval in a sample of individuals with stroke aphasia and age-matched controls.

Theories of brain organization posit that cognitive functions like language are organized in widespread networks consisting of specialized brain regions and their interconnecting white matter fiber tracts (e.g., Hickok and Poeppel, 2007; Saur *et al.*, 2008). *Dual-stream models of language function* (Hickok and Poeppel, 2007; Roelofs, 2014; Saur *et al.*, 2008) assume that language consists of two cortical interface streams: A ventral semantic stream that maps sound to meaning and a dorsal phonological stream that maps sound to articulation. The ventral stream has a weak left-hemisphere bias and projects ventrolaterally to connect sensory speech input to bilateral middle and inferior temporal cortex (Hickok and Poeppel, 2007). The dorsal stream is more strongly left-hemisphere dominant and projects dorsoparietally to the posterior frontal lobe and the posterior-dorsal temporal lobe. Critically, these processing streams are bi-directional and thus support both auditory comprehension (e.g., Hickok and Poeppel, 2007) and speech production (Hula *et al.*, 2020).

The precise white matter substrates of the dual-stream model of language function continue to be debated (Hula *et al.*, 2020), but general agreement is emerging regarding the involvement of certain tracts. The arcuate fasciculus (AF), which participates in phonological and motor-speech processing, is the primary dorsal stream tract (Fernández-Miranda *et al.*, 2015; Glasser and Rilling, 2008; Roelofs, 2014; Saur *et al.*, 2008). Specific portions of the AF may also contribute to semantic processing in language production (Fernández-Miranda *et al.*, 2015; Glasser and Rilling, 2008; Hula *et al.*, 2020; Roelofs, 2014). Semantic processing in the ventral stream is associated with the inferior fronto-occipital fasciculus (IFOF), the uncinate fasciculus (UF), and the extreme capsule (de Champfleur *et al.*, 2013; Han *et al.*, 2013; Hula *et al.*, 2020; Saur *et al.*, 2008). The middle longitudinal fasciculus (MdLF) has anatomically and functionally been associated with both dorsal and ventral streams (e.g., Saur *et al.*, 2008).

Recent work has refined and expanded our understanding of the white-matter pathways that support language, particularly object naming. Hula and colleagues (2020) examined the white matter components of dual-stream language models by investigating structural white matter connectivity associated with semantic and phonological object naming in aphasia. The SP interactive two-step model, a computational model of word production (Dell *et al.*, 2013), was used to derive s and p parameter values from participants’ performance on the Philadelphia Naming Test (Roach *et al.*, 1996) as indices of their semantic and phonological ability. Diffusion spectrum imaging was acquired for each participant, and connectometry analyses (Yeh *et al.*, 2016) examined two multiple regression models to identify the local connectomes associated with the s and p parameters. The results indicated that the structural integrity of both dorsal (AF, MdLF) and ventral tracts (IFOF, UF) was associated with semantic ability, whereas only the integrity of dorsal tracts (AF, MdLF) was associated with phonological ability. In addition, limbic tracts such as the posterior cingulum and fornix were associated with both semantic and phonological processing ability. These results are largely consistent with dual-stream models of language function, but critically indicate that: semantic processing is associated with both dorsal and ventral pathways, the AF is involved in both semantic and phonological processing, and subcortical and limbic structures may also be associated with object naming ability.

A notable limitation of Hula and colleagues’ study (2020) is that they only investigated individuals with aphasia. This is problematic for at least two reasons. First, high lesion overlap in the posterior inferior frontal lobe limited the power to detect behavioral associations with fiber tracts in this region. Second, it is unclear whether the identified tracts are associated with object naming only in response to the aphasia-producing lesion or to what extent the findings reflect tracts associated with naming in healthy, pre-stroke brain organization. Furthermore, Hula and colleagues only examined object naming, not action naming. These shortcomings limit the strength of the conclusions regarding brain-language relationships that may be drawn based on these findings.

In contrast to object naming, there have been fewer investigations of the grey or white matter networks subserving action naming. Extending dual-stream models of language function to action naming would predict that a comparable set of grey and white-matter structures support both object and action naming performance. Conceptual representations, which are likely broadly cortically distributed, would serve as input to dorsal and ventral streams (Hickok and Poeppel, 2007; Saur *et al.*, 2008). These perisylvian cortical regions and associated white-matter tracts would in turn support retrieval of linguistic labels for both objects (nouns) and actions (verbs). These white-matter tracts include IFOF and UF (ventral stream tracts) and AF and MdLF (dorsal stream tracts). Extending dual-stream models in this way would be aligned with traditional *cognitive-linguistic accounts* of language performance and language impairments in aphasia. These theoretical and clinical approaches assume that cognition is a modular system that maintains linguistic representations separate from conceptual information (Goodgalss, 1993; McNeil and Pratt, 2001).

However, there is both theoretical and empirical reason to expect that action naming (verb retrieval) may engage somewhat different neurocognitive systems from those implicated in object naming (noun retrieval). As noted above, verbs and nouns serve different functions in language and can be differentially impaired in aphasia (e.g., Caramazza and Hillis, 1991; Miceli *et al.*, 1988). They also engage distinct types of conceptual knowledge (e.g., Vigliocco *et al.*, 2011). Nouns typically refer to object concepts that are processed by occipito-temporal processing streams (Damasio, 1989), whereas verbs commonly refer to action concepts that are processed via frontoparietal systems (Tranel *et al.*, 2001; Wurm and Caramazza, 2019).

The potential contribution of frontoparietal systems, especially motor regions, to action concepts and action naming is consistent with *grounded-cognition accounts* of action- and verb-knowledge (Tranel *et al.*, 2001; Pulvermüller, 2018). Neuroimaging and neuropsychological evidence suggests that primary motor and premotor cortex are not only activated during verb processing (Hauk *et al.*, 2004; Pulvermüller *et al.*, 2005) but may causally contribute to action naming. For example, patients with neurodegenerative diseases affecting primary motor and premotor cortex have shown correlations between degree of atrophy and impairments of both action knowledge and verb retrieval (e.g. Bak, 2013; Roberts *et al.*, 2017), lesion to pre-motor cortex has predicted both action- and verb-processing impairments post-stroke (Kemmerer *et al.*, 2012), and neurostimulation to motor and pre-motor regions can facilitate (Pulvermüller *et al.*, 2005) or disrupt action-related semantic processing (Gerfo *et al.*, 2008) and can facilitate novel action-word learning (Liuzzi *et al.*, 2010). This evidence is consistent with grounded-cognition models that claim that both language and conceptual processing are rooted in experience-driven, multimodal neural representations that simulate sensory, motor, and introspective states (e.g., Barsalou, 2008).

A limited number of studies have addressed the white-matter substrates of action-naming. The findings of these studies provide support for the predictions of both cognitive-linguistic and grounded-cognition accounts. First, Bello and colleagues reported action naming findings from intraoperative language mapping experiments in awake surgery patients (e.g. Bello *et al.*, 2008). They found that verb processing involved portions of the superior longitudinal fasciculus (SLF), AF, UF, and IFOF. Phonemic paraphasias (word or non-word substitutes that share resemblance to the target word sounds; e.g., dog, gog) were elicited by stimulations to dorsal pathway fiber tracts: SLF, AF, and the subcallosal fasciculus. In contrast, semantic paraphasias (word substitutes related to the target word in meaning; e.g, dog, cat) were produced during stimulation of ventral pathway tracts: UF, IFOF, and ILF. These observations are consistent with dual-stream models of language. However, Bello and colleagues did not systematically stimulate white-matter tracts outside the traditional dorsal and ventral streams. These findings thus do not address whether white-matter structures outside those associated with the dorsal and ventral language streams also critically support action naming.

Second, Akinina and colleagues (2019) recently examined gray and white matter substrates of action naming, focusing in particular on lexical-semantic aspects of action naming. Russian speakers with stroke-induced aphasia and/or dysarthria named black-and-white line drawings of actions. Akinina and colleagues (2019) scored naming performance and categorized errors as most likely reflecting lexical-semantic versus phonological deficits. The frequency of lexical-semantic errors was used as a measure of semantic processing during action naming, like the s-parameter used in Hula and colleagues’ (2020) object naming study. Voxel-based lesion-symptom mapping then examined the relationship between gray and white matter intactness and lexical-semantic error rates, covarying phonological error rate. The analysis revealed that many white matter fibers were associated with lexical-semantic processes of action naming, including the frontal aslant tract (FAT), IFOF, SLF II and III, UF, long segment of the AF, fronto-orbital polar and frontal inferior longitudinal tracts, and the fronto-insular tract. The analysis also highlighted projection fibers that couple the cortex with subcortical regions and the spinal cord, including anterior thalamic projections, corticospinal, frontostriatal and frontopontine tracts.

As previously discussed, the IFOF, UF, and AF are often implicated in object naming and are consistent with dual-stream models of language function. However, the SLF, FAT, and projection fibers are not. These white matter tracts might uniquely subserve action naming because of their roles in action processing and motor control. The SLF, which connects frontal regions to temporo-parietal regions, has been implicated in motor-speech articulation (Duffau *et al.*, 2014), ideomotor apraxia (e.g., Leiguarda and Marsden, 2000), motor function sequences, semantic action processing, and motor imagery (Parlatini *et al.*, 2017). The FAT, which bridges supplementary and pre-supplementary motor areas from the superior frontal gyrus to posterior portions of the inferior frontal gyrus, has been associated with verbal fluency and motor speech initiation (Kinoshita *et al.*, 2015; Kronfeld-Duenias *et al.*, 2016; Sierpowska *et al.*, 2015; Vassal *et al.*, 2014). Finally, projection fibers include anterior thalamic projections that are associated with spatial navigation and memory (e.g., Jankowski *et al.*, 2013), corticospinal tract (CST) projections that are involved in voluntary motor control (Cho *et al.*, 2007; Welniarz *et al.*, 2017; Zhu *et al.*, 2010), frontostriatal circuits that modulate motor control and executive function (e.g,, Morris *et al.*, 2016), and frontopontine tracts that connect cortex to the opposite cerebellum for the coordination of planned motor functions. Akinina and colleagues’ finding that these tracts were associated with lexical-semantic aspects of action naming suggests that a broad set of multimodal connections support verb processing and is consistent with grounded accounts of cognition.

Although these findings are promising, they are limited in several respects. First, as in Hula and colleagues’ (2020) study of object naming, Akinina and colleagues’ (2019) participant sample did not include any adults without lesions. This limits our ability to draw conclusions regarding whether these findings (suggesting that motor networks support action naming) also apply to the operation of the undamaged system, or instead reflect adaptation or reorganization post-stroke. Second, the overlay analysis method that Akinina and colleagues (2019) employed was limited to detecting intersections between portions of tract probability maps and was not capable of tracking the course of specific fibers. This approach therefore results in less stable and reliable estimates of individual white-matter tracts as well as the whole-brain white-matter connectome than the diffusion MRI connectometry methods used in the current study (Yeh *et al.,* 2016). As a result, this approach is less able to estimate the strength of the relationships between either specific tract integrity or individuals’ white-matter connectome and action-naming performance.

The current study employs diffusion MRI connectometry (Yeh *et al.*, 2016), the same analytic approach followed by Hula and colleagues (2020), to examine the specific white matter components of *action naming* networks rather than object naming networks. Connectometry uses correlational tractography, a new tractography modality that tracks the exact segment of pathways with anisotropy correlated with the study variable. It has greater sensitivity than conventional voxel-based or tract-based analyses (Yeh *et al.*, 2016). Moreover, we utilized beyond-tensor diffusion MRI acquisition to address the limitation of conventional diffusion MRI tensor analysis (Yeh *et al.*, 2013). These methods provide a superior estimate of the white-matter tracts supporting action naming to voxel-based white-matter overlay methods (e.g., Akinina et al., 2019). Further improving on Hula and colleagues (2020) and Akinina and colleagues (2019), the current study tested both individuals with aphasia and healthy control participants without stroke aphasia. This enables stronger conclusions to be drawn regarding how different neural systems support language performance. The connectometry analyses tested two contrasting sets of theoretical predictions regarding verb retrieval, namely that it is associated with the integrity of: (1) both motor and language tracts, as predicted by grounded cognition accounts, or (2) only language tracts, as predicted by cognitive-linguistic accounts. Our specific hypotheses follow from our literature review, which identified tracts implicated in dual-stream language models (AF, MdLF, IFOF, UF) *or* predominantly motor pathways (FAT, SLF, CST, frontopontine tract, other projection tracts).

## 2. Materials and Methods

### 2.1. Participants

Participants were 14 individuals with chronic aphasia due to unilateral left-hemisphere stroke and 15 age-matched neurotypical controls. All participants were native English speakers, able to provide informed consent, 25-85 years old, (pre-morbidly) right-handed, and had no history of progressive neurological or psychiatric disease, drug/alcohol dependence, or significant mood or behavioral disorder. Individuals with aphasia were more than 6 months post-onset (range: 19-265 months; M=88.9, SD=66.5 months), had a Comprehensive Aphasia Test (CAT; Swinburn *et al.*, 2004) Naming Modality T-score ≥40, and an overall mean T-score <70. Participant demographics are reported in Table 1 for participants with aphasia and Table 2 for controls.

**Table 1.**
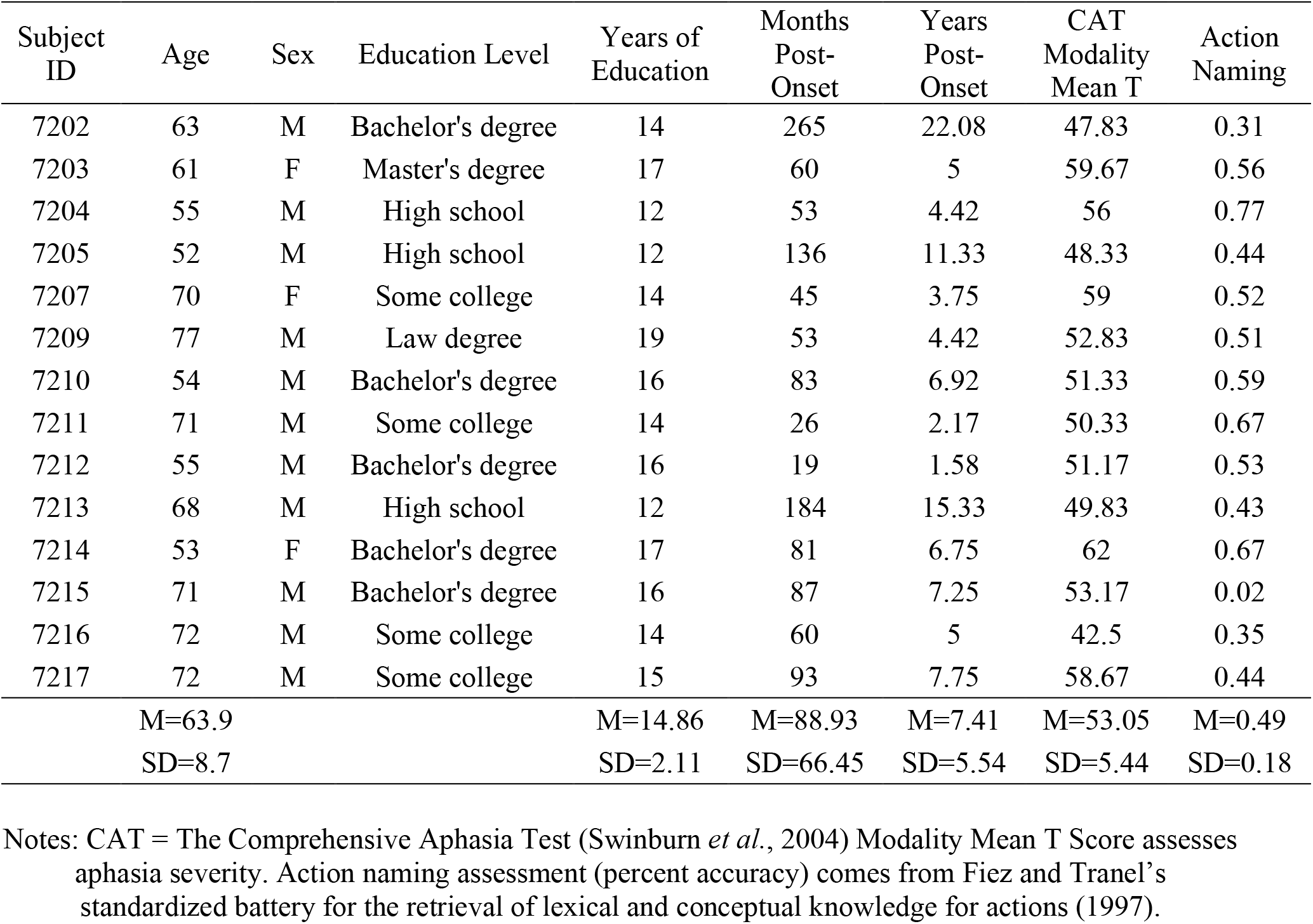
Demographic characteristics of participants with aphasia.

**Table 2.** Demographic characteristics of age-matched control participants.

| Participant ID | Age | Sex | Education Level | Years of Education | Action Naming |
| --- | --- | --- | --- | --- | --- |
| 7001 | 42 | M | Tech college | 14.5 | 0.90 |
| 7002 | 59 | M | High school | 12 | 0.99 |
| 7003 | 74 | M | Bachelor's degree | 16 | 0.90 |
| 7004 | 52 | M | Bachelor's degree | 16 | 0.86 |
| 7005 | 54 | M | Bachelor's degree | 16 | 0.94 |
| 7006 | 57 | M | High school | 12 | 0.80 |
| 7007 | 72 | F | Master's degree | 18 | 0.92 |
| 7008 | 64 | F | Master's degree | 18 | 0.95 |
| 7010 | 74 | M | Master's degree | 20 | 0.92 |
| 7011 | 68 | F | Master's degree | 22 | 0.98 |
| 7012 | 72 | M | Bachelor's degree | 16 | 0.95 |
| 7013 | 65 | M | Law degree | 19 | 0.87 |
| 7014 | 71 | F | Master's degree | 17 | 0.93 |
| 7015 | 52 | M | Master's degree | 22 | 0.94 |
| 7016 | 69 | F | Master's degree | 18 | 0.89 |
| Summary | M=63<br>SD=9.82 | 5 F; 10 M |  | M=17.1<br>SD=3 | M=0.92<br>SD=0.05 |

Notes: Action naming assessment (percent accuracy) comes from Fiez and Tranel's standardized battery for the retrieval of lexical and conceptual knowledge for actions (1997).

Participants who met any of the following criteria were excluded: 1) significant hearing loss or vision impairment that prevented experiment completion, or 2) pre-existing or subsequent brain injury/stroke (e.g., to right-hemisphere regions for individuals with aphasia). Participants were excluded if their T scores were less than 30 for the CAT Cognitive Screening semantic memory or recognition memory subtests. T scores under 30 would be indicative of frank auditory, visual, motor speech, or general cognitive deficits. In addition, neurotypical participants were excluded if they failed a line-bisection visual screening, a binaural pure-tone hearing screening (0.5, 1, 2, and 4 KHz at 40 dB), a Mini-Mental State Examination (required 27/30; Folstein *et al.*, 1975), or Raven’s Coloured Progressive Matrices (required 30/36; Raven, 1965).

Each participant provided written informed consent to participate and authorization for release and review of relevant medical records for research purposes. Participants were compensated for their time participating in both behavioral and neuroimaging protocols.

### 2.2. Behavioral language materials and procedures

All participants were administered a standardized action naming task (Fiez and Tranel, 1997) that consists of 100 colored photographs of actions drawn from a variety of semantic categories. See Fiez and Tranel (1997) for task details. The action naming task was administered and scored according to standardized procedures (Fiez and Tranel, 1997; Kemmerer *et al.*, 2001, 2012). Participants were given detailed instructions and practice items, to ensure they understood the task. For each item, a computer displayed a colored photograph of a person or animal performing an action. The participant was instructed to provide a single verb that described what the person or animal was doing. Participants were allowed an unlimited amount of time to name each action.

An external microphone recorded naming responses in Audacity^®^ software, and trial-level accuracy was scored by hand for each task. The participant’s first response was recorded. Following Fiez and Tranel (1997), experimenters prompted participants for a second response if the participant did not provide a verb (“Can you tell me what the person is *doing*?”) or if the participant provided a description (“Can you give me a *single word* that best describes what the person is doing?”). Target responses provided after a prompt were scored as correct. Alternate forms of a target verb were also accepted as correct (e.g. run, running, ran). Normative data from Fiez and Tranel (1997) reported 85 percent average accuracy, with a standard deviation of 5 percent.

### 2.3. MRI data acquisition and reconstruction

Diffusion spectrum imaging (DSI) scans were acquired on a 3T SIEMENS Tim Trio scanner using a 257-direction 2D EPI diffusion sequence (TE=150 ms, TR=3439 ms, voxel size=2.4×2.4×2.4 mm, FoV=231×231 mm, b-max=7000 s/mm2). A diffusion sampling length ratio of 1.25 was used, and the output resolution was 2 mm isotropic. The restricted diffusion was quantified using restricted diffusion imaging. The diffusion data were reconstructed in the MNI space using q-space diffeomorphic reconstruction to obtain the spin distribution function. The anisotropy values were extracted from the data and used in the connectometry analysis to derive correlational tractography. T1 and T2-weighted images were acquired, lesion masks were constructed, and lesion volumes were calculated for each participant.

### 2.4. Data analysis

Diffusion MRI connectometry (Yeh *et al.*, 2016) analyses were conducted in DSI Studio (http://dsi-studio.labsolver.org). All 29 participants’ diffusion MRI scans were included in a connectometry database. A multiple regression model was used to identify the local connectome associated with action naming performance, using a deterministic fiber tracking algorithm (Yeh *et al.*, 2013) with an assigned T-score threshold of 3.00 and a fiber length threshold of 30 mm. Lesion volume was included as a covariate, with zero entered for control participants. Topology-informed pruning was conducted to remove false connections. All tracks generated from bootstrap resampling were included. The seeding number of each permutation was 50,000. To estimate the false discovery rate, a total of 2,000 randomized permutations were applied to obtain the null distribution of the track length.

## 3. Results

White matter tracts identified as *positively* associated (false discovery rate [FDR]=0.000071) with action naming include portions of the left corticothalamic pathway (37%), left IFOF (17%), corpus callosum (12%), left frontopontine tract (6.1%), left cingulum (5.1%), left parietopontine tract (4.4%), left ILF (3.9%), left AF (3.4%), left corticostriatal pathway (3%), left CST (2%), right corticothalamic pathway (1.5%), anterior commissure (1.2%). See Figures 1-2. Percentages reflect the proportion of total streamlines identified in the analysis that were associated with each tract. Figure 3 shows results from the sagittal view, color coded for specific tract assignment. These results identify tracts associated with neurocognitive dual-stream models of language function (Figure 4) as well as projection pathways that connect frontal and parietal cortices to subcortical regions associated with motor programming (Figure 5). Approximately 75 percent of the tracts associated with verb retrieval were projection pathways. The analysis showed no white matter tracts whose connectivity was reliably *negatively* associated with action naming performance (FDR=1). False fibers were not interpreted.

**Figure 1.**
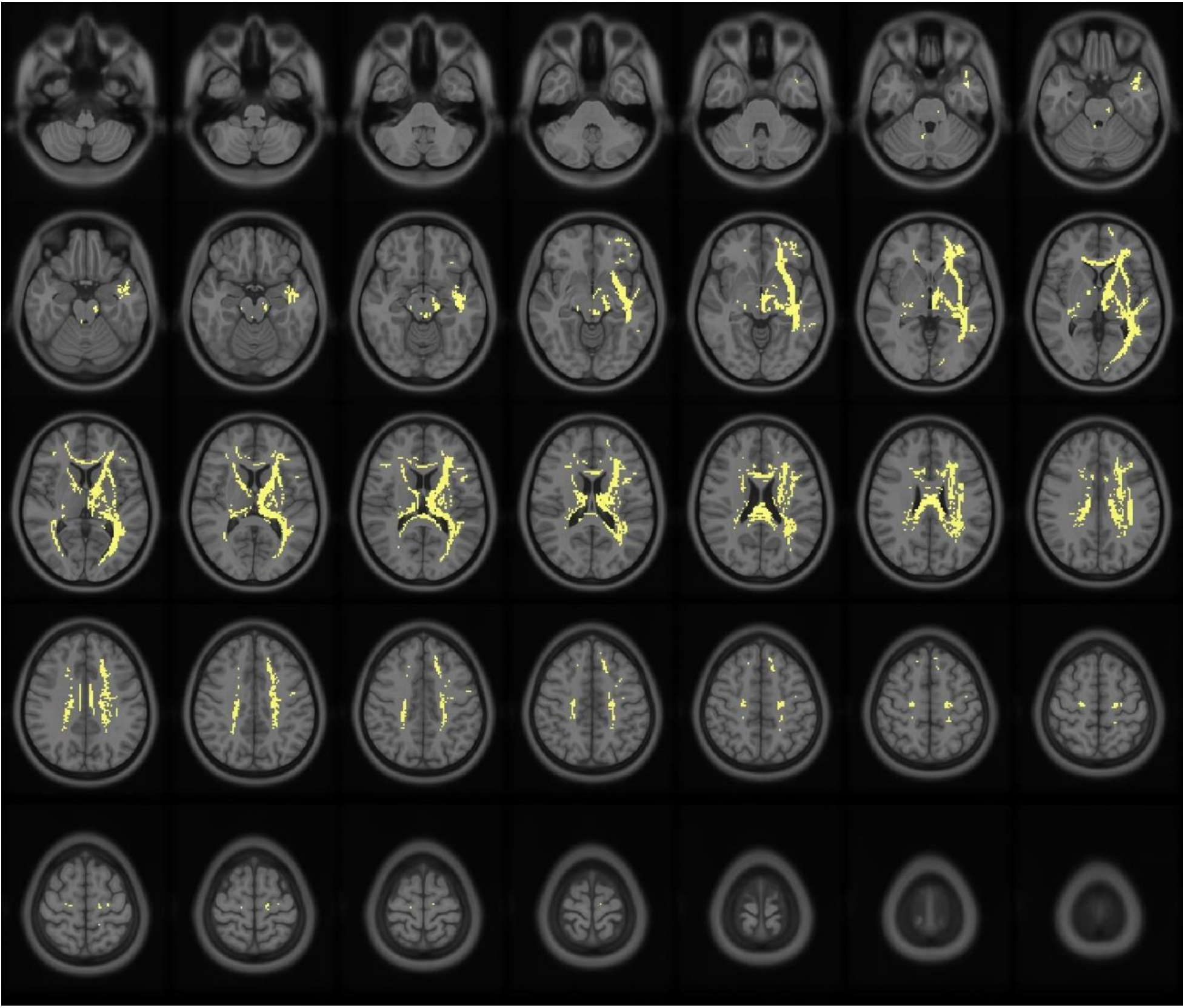
Voxel mapping of white matter fiber tracks with quantitative anisotropy positively correlated with verb naming in all participants.

**Figure 2.**
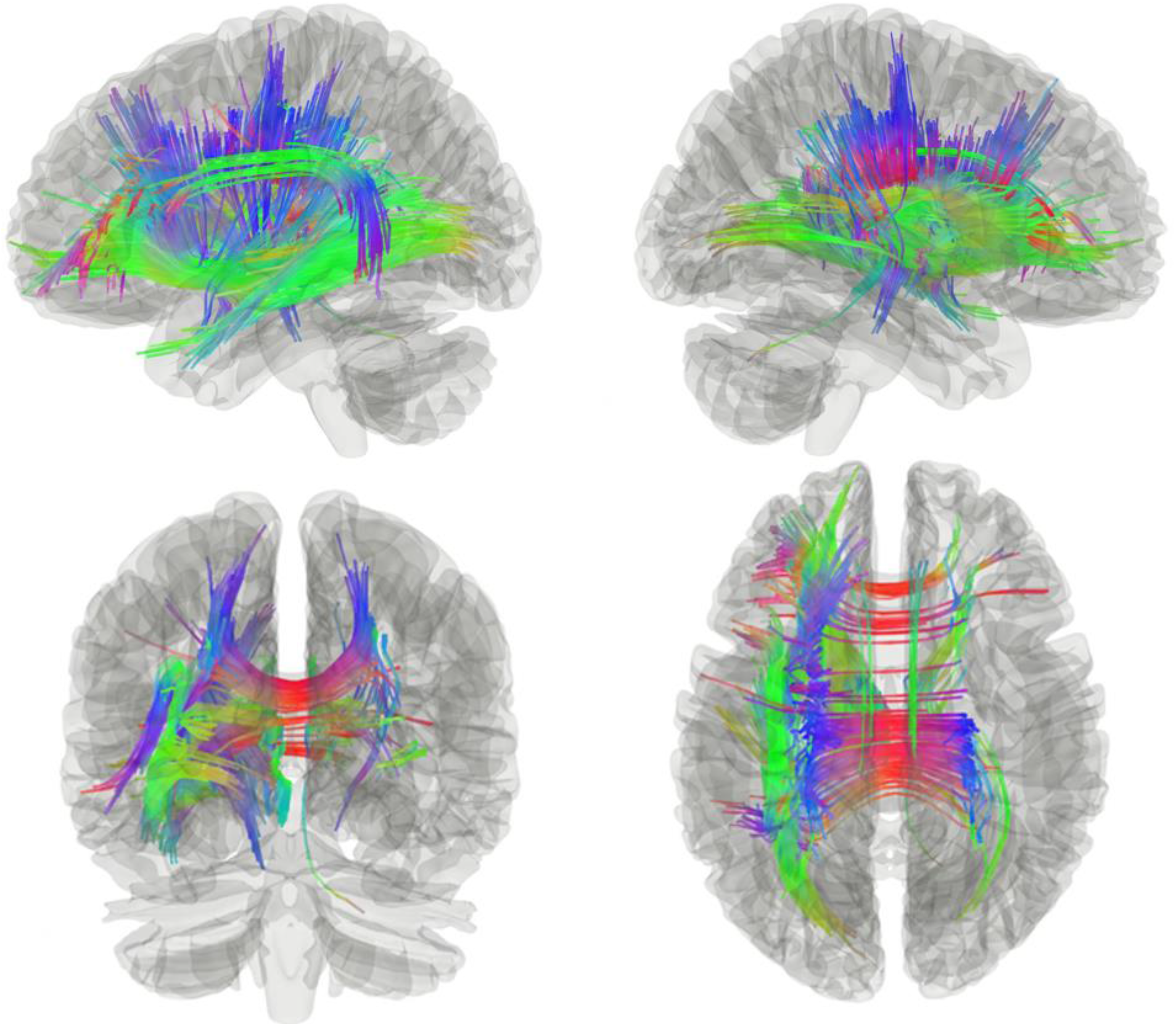
White matter fiber tracks positively correlated with verb naming in all participants. Notes: Colors reflect fiber orientation. Blue: superior–inferior. Green: anterior–posterior. Red: medial– lateral.

**Figure 3.**
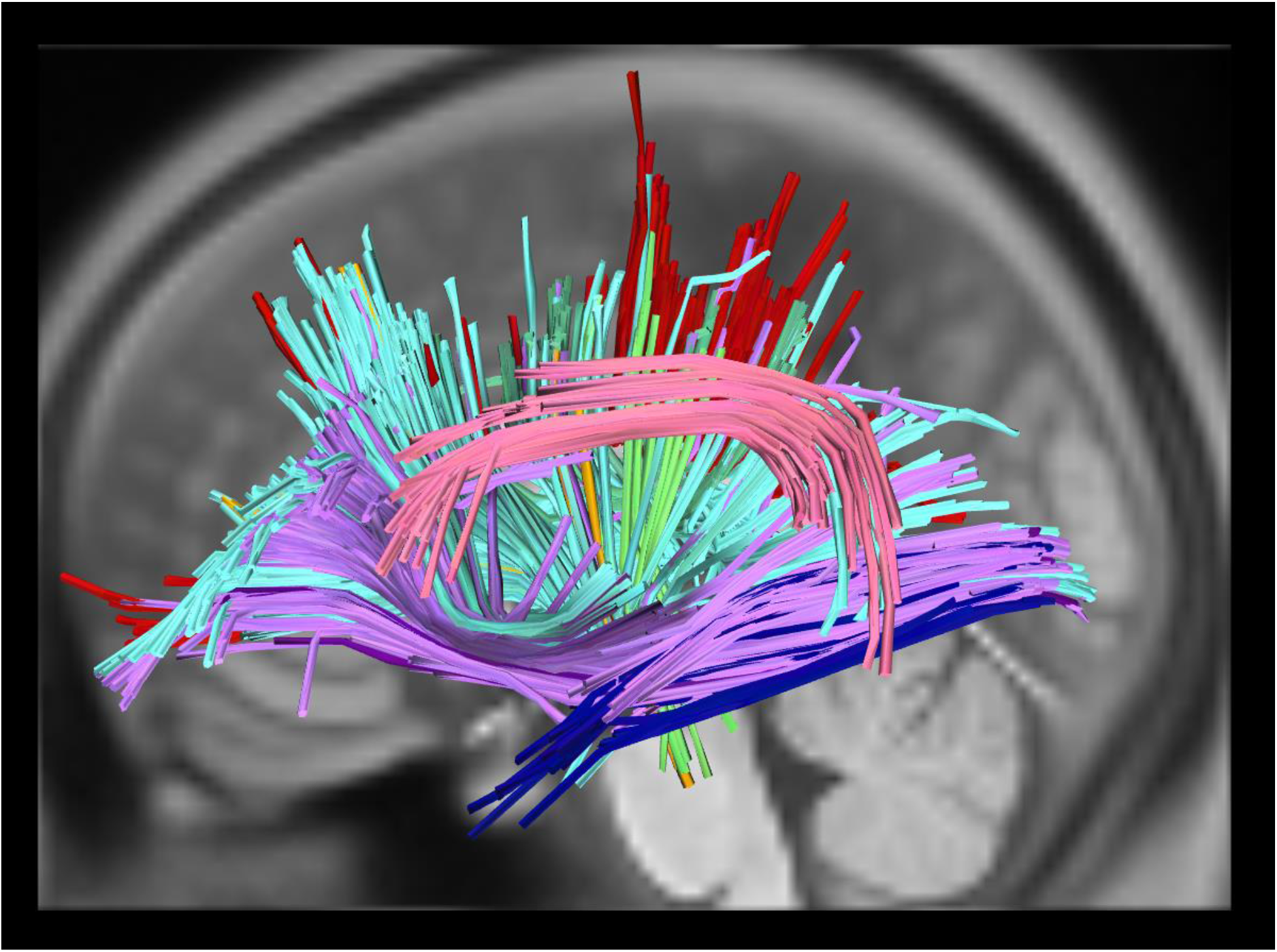
White matter streamlines associated with each tract. Notes: Colors reflect tract assignment. Red – Corpus callosum; Orange – Left frontopontine tract; Pink – Left arcuate fasciculus (AF); Lime green – Left parietopontine tract; Forest green – Left cingulum; Pale blue – Left corticothalamic pathway; Navy – Left inferior longitudinal fasciculus (ILF); Lilac – Left corticostriatal pathway; Violet – Left inferior frontooccipital fasciculus (IFOF); Gray – Left corticospinal tract (CST).

**Figure 4.**
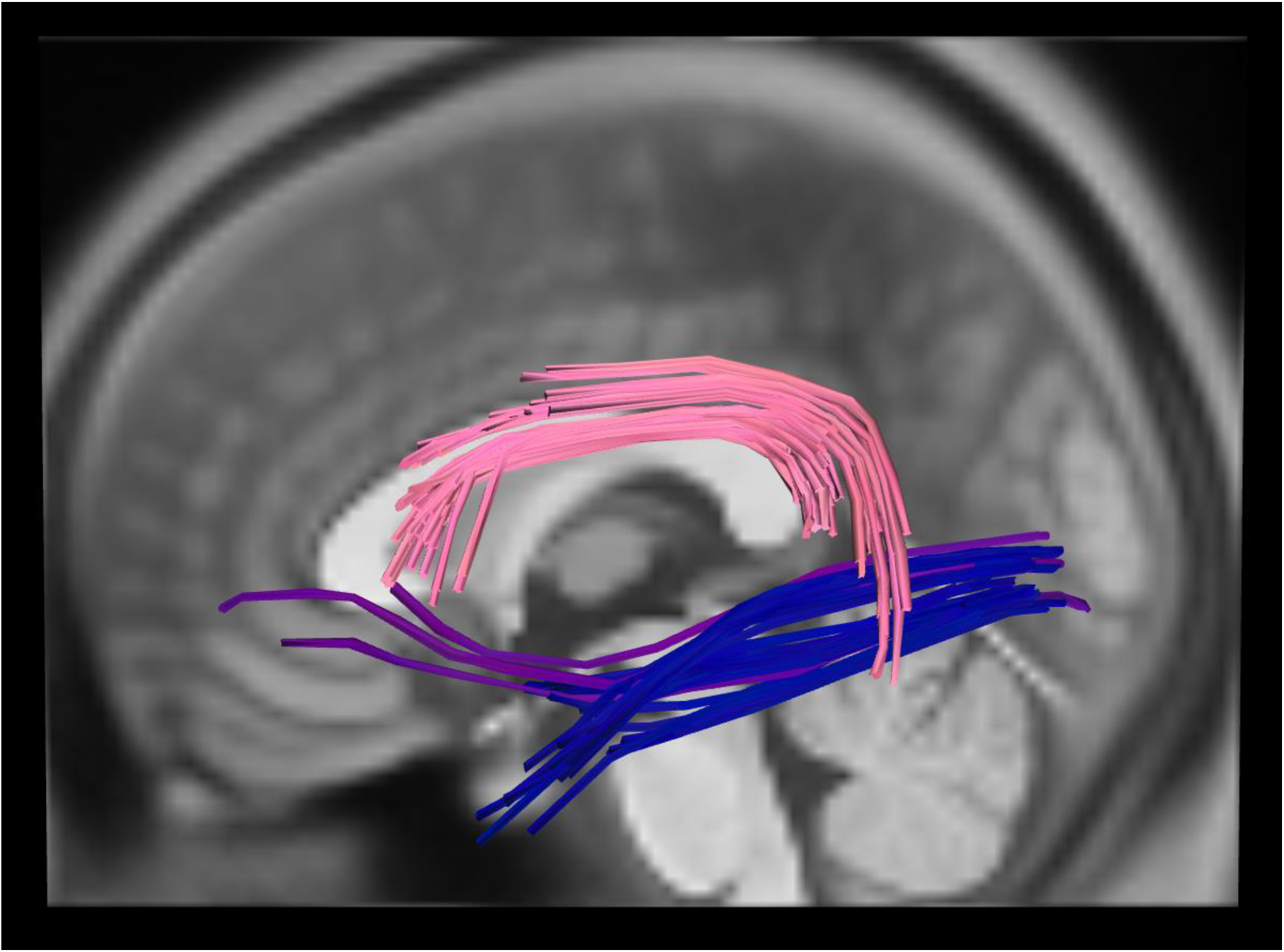
Identified white matter tracts associated with dual-stream models of language function. Notes: Colors reflect tract assignment. *Dorsal stream:* Pink – Left arcuate fasciculus (AF). *Ventral stream:* Navy – Left inferior longitudinal fasciculus (ILF); Violet – Left inferior frontooccipital fasciculus (IFOF).

**Figure 5.**
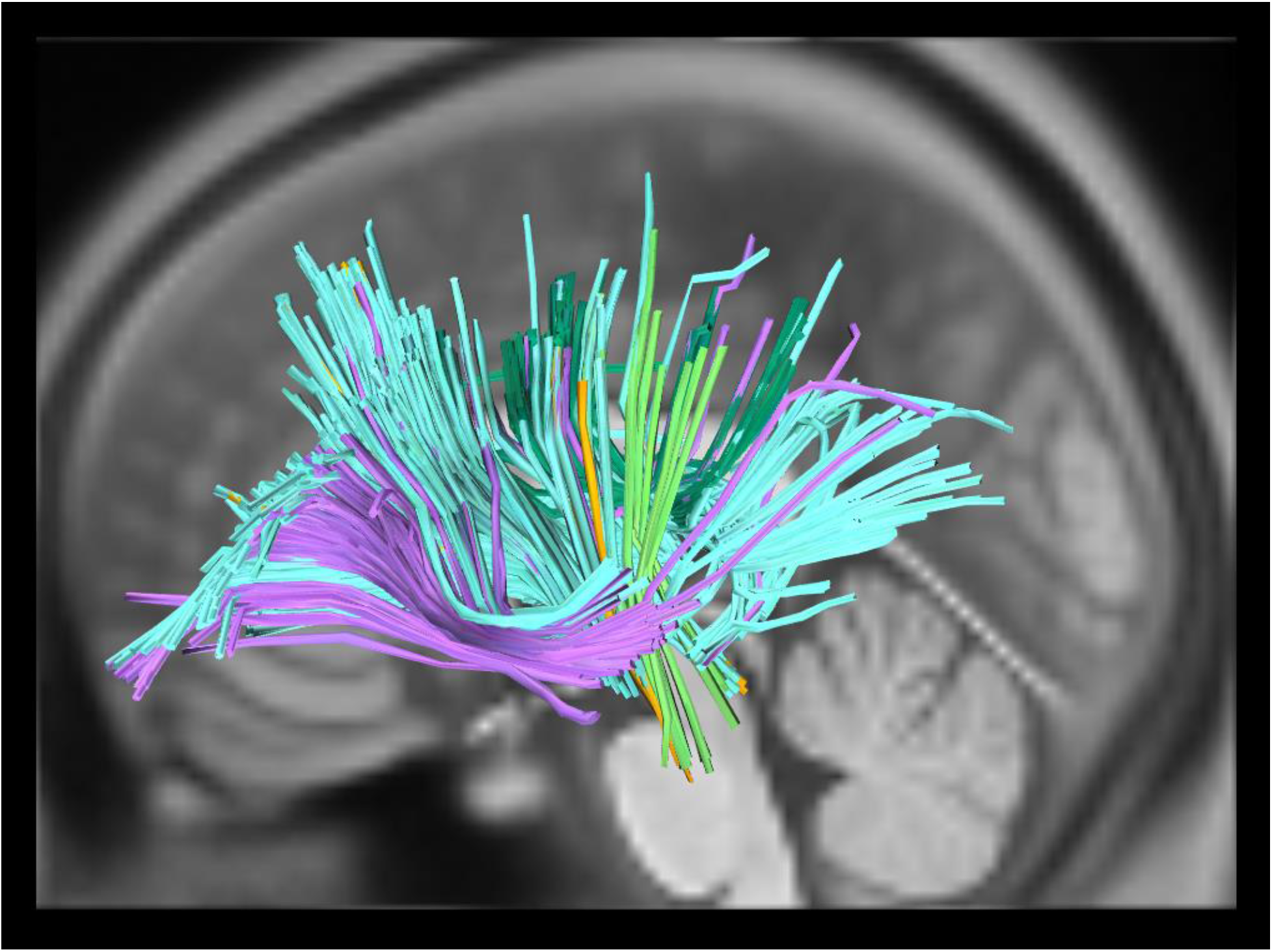
Identified white matter tracts associated with projection motor pathways. Notes: Colors reflect tract assignment. Orange – Left frontopontine tract; Lime green – Left parietopontine tract; Forest green – Left cingulum; Pale blue – Left corticothalamic pathway; Lilac – Left corticostriatal pathway; Gray – Left corticospinal tract (CST).

As expected given the left-hemisphere lateralization of language and patient inclusion criterion of unilateral left-hemisphere stroke, the positive association between action naming and white-matter tract integrity was more prominent in the left than in the right hemisphere. However, the corpus callosum and several right hemisphere tracts were also robustly associated with action naming, suggesting bilateral neural involvement in verb retrieval (Figures 1-2).

## 4. Discussion

This study identified white matter tracts associated with verb retrieval (action naming), using diffusion spectrum MRI connectometry in 14 participants with aphasia due to left-hemisphere stroke and 15 age-matched controls. Previous work characterized white-matter correlates of noun-production deficits (Hula *et al.*, 2020) and found results largely consistent with dual-stream language models. Cognitive-linguistic theories predict that the same dorsal and ventral pathways that underlie noun production should underlie (respectively, phonological and lexical-semantic aspects of) verb production. In contrast, grounded cognition accounts predict that distributed pathways supporting both linguistic and non-linguistic conceptual-motor performance critically support action naming.

Verb production and verb-retrieval deficits were assessed by a 100-item standardized task of action picture naming (Fiez and Tranel, 1997). The degree of connectivity between adjacent voxels was used to reconstruct a structural connectome for each participant (Yeh *et al.*, 2016). Verb production ability was then regressed on a connectome matrix for the full group of participants, controlling for lesion volume. This approach improves upon methods that map pathways between cortical origin and termination points by providing more specific and reliable connectome reconstruction. In comparison, the overlay analysis method for white matter that Akinina and colleagues (2019) employed was limited to detecting intersections between portions of tract probability maps and was not capable of tracking the course of specific fibers.

Connectometry analyses identified tracts associated with verb retrieval that were largely consistent with predictions of grounded cognition accounts: Connectivity of classic linguistic *and* motor pathways were positively associated with verb retrieval ability. First, verb retrieval was positively associated with the integrity of several linguistic tracts, including left AF, IFOF, and ILF. These results are consistent with dual-stream models of language function and observations regarding white matter correlates of object naming (e.g., Han *et al.*, 2013; Hula *et al.*, 2020), action naming (Akinina *et al.*, 2019), and verb processing in intraoperative stimulation experiments (Bello *et al.*, 2008). The left IFOF and ILF have consistently been associated with the ventral stream and the process of mapping sound to meaning (e.g., Han *et al.*, 2013; Hula *et al.*, 2020; Saur *et al.*, 2008). The left AF is considered the predominant dorsal tract in the dual-stream language models, but this was the only dorsal language tract robustly associated with verb retrieval.

The MdLF and UF have both been characterized as language tracts previously but were not identified in the current analyses. The MdLF has been implicated in tractographic studies of language processing (de Champfleur *et al.*, 2013) and has been associated with both semantic and phonological stages of object naming in adults with aphasia (Hula *et al.*, 2020). However, evidence from patients with resected MdLF suggests that this tract is not essential for language processing (Hamer *et al.*, 2011). Similarly, the UF was not identified in our analyses, but has been characterized as a ventral-stream tract associated with language function and semantic processing (Han *et al.*, 2013), including semantic paraphasias in object naming (Hula *et al.*, 2020) and action naming (Bello *et al.*, 2008; Akinina *et al.*, 2019). However, there is also evidence that the UF is not systematically essential for language and that alternative, perhaps indirect ventral pathways may functionally compensate when the UF is damaged or partially resected (Duffau *et al.*, 2009). This possibility and the specific contributions of the MdLF and UF to verb retrieval and broader language function must be explored further.

Second, verb-retrieval deficits were associated with compromised integrity to several motor pathways, including bilateral corticothalamic tracts, and left corticospinal, corticostriatal, frontopontine, and parietopontine tracts. Although these tracts are not traditionally associated with language, frontostriatal, frontopontine, and anterior thalamic projections have been associated with lexical-semantic stages of action naming in a VLSM analysis of individuals with aphasia (Akinina *et al.*, 2019). The current results identified positive associations between verb retrieval and tract integrity connecting both frontal and parietal regions of cortex to the striatum of the basal ganglia, as well as to the thalamus. In particular, the majority of the corticostriatal tracts identified connected left hemisphere cortex to the left putamen, which has been previously associated with movement disorders like Tourette syndrome and Parkinson’s disease (e.g., DeLong and Wichmann, 2007), as well as motor functions including motor planning, learning, and execution (e.g., Watkins and Jenkinson, 2016). In addition, both Akinina and colleagues (2019) and the current results identified the involvement of corticospinal and corticopontine tracts. As discussed in the introduction, there is a substantial body of work linking this set of projection fibers to voluntary motor control and action-related cognition. The involvement of these motor pathways in verb retrieval is therefore in line with predictions of grounded cognition accounts.

It is unclear why the FAT and SLF were not associated with verb retrieval here, as they were in Akinina *et al.* (2019), but these tracts are not typically identified in picture naming experiments. In addition, the pars opercularis of the inferior frontal gyrus (BA 44) hosts terminations of both the FAT and the SLF III (as well as the AF; Rojkova *et al.*, 2016). This region has a high probability of being damaged in individuals with stroke aphasia and the statistical power to detect meaningful relationships between these tracts and naming performance may therefore be limited. Regardless, it is worth noting that although the result was not robust, approximately 500 fibers of the left SLF were associated with verb naming. The stronger SLF and FAT relationships identified by Akinina and colleagues (2019) could reflect a wider range of articulation and verbal fluency among participants in that study (*articulation/SLF:* Duffau *et al.*, 2014; *fluency/FAT:* Kinoshita *et al.*, 2015; Kronfeld-Duenias *et al.*, 2016; Vassal *et al.*, 2014). As Akinina and colleagues (2019) discuss, fluency impairments may be partially related to word-retrieval deficits in their sample, and further investigation is needed to characterize white matter tracts associated with fluency’s role in word retrieval. If this speculation is correct, it suggests that Akinina and colleagues’ results reflected the contribution of the FAT to verbal fluency rather than to verb retrieval. However, Kemmerer and colleagues (2012) also did not find associations between verb retrieval and SLF or FAT in a sample of adults with a wide variety of lesion locations. Future work should continue to assess the roles of these tracts in word retrieval, articulation, and fluency in large samples of diverse patients.

Finally, the corpus callosum and cingulum were also robustly associated with verb retrieval. Hula and colleagues (2020) similarly found the splenium of the corpus callosum was positively associated with phonological and semantic abilities during object naming. The present study also found associations with bilateral cinguli, although more left than right hemisphere tracts. Cingulum bundles have been related to many functions, including emotion, motivation, executive function, spatial processing, and motor function (Bubb *et al.*, 2018). Bilateral cinguli were associated with both phonological and semantic function in object naming – with an antero-posterior distinction between the two, respectively (Hula *et al.*, 2020). The involvement of the corpus callosum and bilateral cinguli is consistent with aphasia models that contend the right hemisphere compensates for stroke-related language impairments (Stefaniak *et al.*, 2020).

A complete discussion of each identified pathway goes beyond the scope of this study. However, these results emphasize that in addition to cortico-cortical networks of dual-stream language models, cortico-subcortical networks, subcortical substrates, and right-hemisphere tracts may be critically involved in action naming and verb-retrieval deficits in aphasia. Furthermore, the present connectometry methods provide new evidence by tracking each fiber tract along its respective course, rather than simply detecting associations with different portions of tract probability maps, as done by Akinina and colleagues (2019).

This study extended connectometry approaches in aphasia to examine the white matter tracts associated with verb-retrieval deficits, and furthermore addressed a critical limitation and threat to external validity by including an age-matched control group. Hula and colleagues (2020) did not include a control group in their connectomtery-based analyses of object naming, nor did Akinina and colleagues (2019) in their VLSM-based analysis of action naming. Including the control group strengthens the conclusions that can be drawn from the current white-matter connectivity findings regarding brain-language relationships. Specifically, it suggests that the role of motor pathways in supporting action naming does not simply reflect reorganization or adaptation post-stroke, but instead holds both for adults with and without lesions to language networks, as assumed by grounded accounts of cognition (Barsalou, 2008; Pulvermüller, 2018). Furthermore, the current findings may help explain a potentially counterintuitive finding reported by Hula and colleagues (2020). Their analyses found robust negative associations between language behavior and fiber tract connectivity, indicating that individuals with lower white-matter integrity (i.e., higher damage) in certain tracts had better language performance. The lack of negative associations in the current study suggest that this previous pattern of findings is likely an artifact due to the sample bias of only including participants with left-hemisphere lesions. Furthermore, the current results are not limited to ambiguous interpretations driven by high lesion overlap and too few tract observations in posterior inferior frontal lobe.

As predicted by grounded cognition accounts, our findings suggest verb retrieval is associated with a distributed network of tracts supporting both linguistic and non-linguistic conceptual-motor performance. These findings are consistent with language operating as a general-purpose cognitive system that engages distributed, multimodal representations across regions specialized for linguistic, motor, and affective/limbic processing (e.g., Barsalou, 2008; Martin, 2016; Zwaan and Madden, 2005). These results are furthermore consonant with previous neuropsychological and neurostimulation studies that found associations between verb/action processing and primary motor, premotor, and supplementary motor cortices (e.g., Kemmerer *et al.*, 2012; Pulvermüller *et al.*, 2005).

## 5. Conclusion

This study was among the first investigate the white matter connectivity associated with action naming abilities and verb-retrieval deficits in chronic left-hemisphere stroke aphasia. The current findings extend beyond prevailing neurocognitive dual-stream models and cognitive-linguistic theories of aphasia by identifying associations between verb retrieval and white matter tracts involved in motor pathways. This new evidence is consonant with grounded accounts of cognition. Furthermore, it underlines the importance of actions and verbs to the development of comprehensive models of brain-language relations.

## Abbreviations

AF: arcuate fasciculus
CAT: Comprehensive Aphasia Test
CST: corticospinal tract
DSI: diffusion spectrum imaging
FAT: frontal aslant tract
FDR: false discover rate
IFOF: inferior fronto-occipital fasciculus
ILF: inferior longitudinal fasciculus
MdLF: middle longitudinal fasciculus
MRI: magnetic resonance imaging
SLF: superior longitudinal fasciculus
UF: uncinate fasciculus
VLSM: voxel-based lesion-symptom mapping

## Author Contributions

H.D., W.H., and M.W.D. conceived of the presented idea. H.D. carried out the experiment. H.D. performed the computations and W.H. and F.Y. verified the analytical methods. H.D. wrote the manuscript with support from W.H., M.W.D, and T.W. All authors discussed the results and contributed to the final manuscript.

## Acknowledgments

The authors gratefully acknowledge the contributions of the High-Definition Fiber Tractography Lab, the Language and Brain Lab, Angela Grzybowski, Alyssa Autenreith, Jessica Barrios, and Yutong Zhang to this work.

